# METTL16 Controls Kaposi’s Sarcoma-Associated Herpesvirus Replication by Regulating *S*-Adenosylmethionine Cycle

**DOI:** 10.1101/2023.06.07.544118

**Authors:** Xinquan Zhang, Wen Meng, Jian Feng, Xinghong Gao, Chao Qin, Pinghui Feng, Yufei Huang, Shou-Jiang Gao

## Abstract

Oncogenic Kaposi’s sarcoma-associated herpesvirus (KSHV) consists of latent and lytic replication phases, both of which are important for the development of KSHV-related cancers. As one of the most abundant RNA modifications, *N^6^*-methyladenosine (m^6^A) and its related complexes regulate KSHV life cycle. However, the role of METTL16, a new RNA methyltransferase, in KSHV life cycle remains unknown. In this study, we have identified a suppressive role of METTL16 in KSHV lytic replication. METTL16 knockdown increased while METTL16 overexpression reduced KSHV lytic replication. METTL16 binding to and writing of m^6^A on MAT2A transcript are essential for its splicing, maturation and expression. As a rate-limiting enzyme in the methionine- *S*-adenosylmethionine (SAM) cycle, MAT2A catalyzes the conversion of L-methionine to SAM required for the transmethylation of protein, DNA and RNA, transamination of polyamines, and transsulfuration of cystathionine. Consequently, knockdown or chemical inhibition of MAT2A reduced intracellular SAM level and enhanced KSHV lytic replication. In contrast, SAM treatment was sufficient to inhibit KSHV lytic replication and reverse the effect of the enhanced KSHV lytic program caused by METTL16 or MAT2A knockdown. Mechanistically, METTL16 or MAT2A knockdown increased while SAM treatment decreased the intracellular reactive oxygen species level by altering glutathione level, which is essential for efficient KSHV lytic replication. These findings demonstrate that METTL16 suppresses KSHV lytic replication by modulating the SAM cycle to maintain intracellular SAM level and redox homeostasis, thus illustrating the linkage of KSHV life cycle with specific m^6^A modifications, and cellular metabolic and oxidative conditions.

## Introduction

Kaposi’s sarcoma-associated herpesvirus (KSHV) is etiologically associated with Kaposi’s sarcoma (KS), primary effusion lymphoma (PEL), multicentric Castleman’s disease (MCD) and KSHV inflammatory cytokine syndrome (KICS) (Damania and Dittmer, 2023; He et al., 2019). The life cycle of KSHV has two phases including latency and lytic replication. During latency, there is no production of virions and only a few viral latent genes including vFLIP, vCyclin, LANA and a cluster of 12 precursor microRNAs located at the latency locus are expressed, which facilitates KSHV evasion of host immune response and persistent infection (Qin et al., 2017; Ye et al., 2011a). Upon stimulation by intracellular or extracellular signals, KSHV can be reactivated from latency into lytic replication, expressing most viral lytic genes and producing infectious virions. The expression of KSHV immediate-early gene, the replication and transcription activator (RTA) encoded by ORF50, is necessary and sufficient for the initiation of viral lytic replication, and expression of downstream lytic genes including early lytic genes such as the mRNA transport and accumulation (MTA) protein encoded by ORF57 and polyadenylated nuclear RNA (PAN), and late lytic genes such as small capsid protein encoded by ORF65 (Ye et al., 2011a). In KS tumors, most of the tumor cells are latently infected by KSHV indicating an essential role of viral latency in the development of KS despite a role of viral lytic replication in the initiation and progression of KS tumors (Ganem, 2010).

*N^6^*-methyladenosine (m^6^A) is the most prevalent mRNA modification and is involved in all aspects of RNA biology including splicing, stability, nuclear export, transcript translation, and formation of high-level structure (Hsu et al., 2017; Jiang et al., 2021; Liu et al., 2015; Wang et al., 2014; Zheng et al., 2013). Deposition of m^6^A on transcripts is primarily catalyzed by a large methyltransferase complex which consists of core proteins METTL3, METTL14 and WTAP, and their associated proteins RBM15, VIRMA, ZC3H13 and HAKAI (Huang et al., 2021). Recent studies have revealed other m^6^A methyltransferase complexes or components including METTL16, and two ribosomal RNA methyltransferases METTL5 and ZCCHC4 which methylate 18S rRNA and eukaryotic 28S rRNA, respectively (Pendleton et al., 2017; Shima et al., 2017; van Tran et al., 2019). METTL16 has been shown to methylate specific transcripts including U6 small nuclear RNA and MAT2A mRNA that encodes *S*-adenosylmethionine (SAM) synthetase (Pendleton et al., 2017; Shima et al., 2017). METTL16 methylates 6 conserved hairpins (hp) in the MAT2A 3’ UTR. Under a low level of SAM, a methyl donor of m^6^A, METTL16 binds to and methylates hp1 to promote MAT2A splicing and hence increases the level of mature MAT2A mRNA expression. However, at a high SAM concentration, METTL16 binding to hp1 site is weakened resulting in the degradation of MAT2A transcript. Therefore, METTL16 behaves as both m^6^A writer and reader for MAT2A transcript to regulate SAM synthetase expression and SAM production (Pendleton et al., 2017).

The m^6^A modifications on viral and cellular RNAs have been profiled during infections of numerous viruses and demonstrated to regulate viral replication (Gokhale et al., 2016; Imam et al., 2018; Kennedy et al., 2016; Lang et al., 2019; Lichinchi et al., 2016a; Lichinchi et al., 2016b; Tan et al., 2018; Tirumuru et al., 2016). Several m^6^A- related proteins including METTL3, FTO, YTHDC1, YTHDF2, RBM15 and SND regulate KSHV replication and the expression of viral genes (Baquero-Perez et al., 2019; Hesser et al., 2018; Martin et al., 2021; Tan et al., 2018; Ye et al., 2017). However, the role of METTL16 in KSHV life cycle has not been examined.

In this study, we have found that METTL16 and METTL16-specific m^6^A modifications control KSHV lytic replication by regulating MAT2A transcript splicing and protein expression to maintain the intracellular SAM level and reduce reactive oxidative stress (ROS). These results reveal a novel role of METTL16 and m^6^A in KSHV life cycle, and offer an insight into the complex network of cellular epitranscriptomics, metabolism and redox homeostasis that regulates KSHV persistent infection.

## Materials and Methods

### Cell culture

RGB-BAC16-infected iSLK (iSLK-RGB-BAC16) cells were cultured in Dulbecco modified Eagle medium (DMEM) supplemented with 10% fetal bovine serum (FBS), 1% penicillin-streptomycin, 1.2 mg/ml hygromycin, 1 µg/ml puromycin, and 250 µg/ml G418 (Brulois et al., 2014). iSLK cells stably express doxycycline (Dox)-inducible RTA (Brulois et al., 2014). The expression of RTA is essential and sufficient for triggering KSHV lytic replication. Primary rat embryonic metanephric mesenchymal precursor cells (MM cells), which are susceptible to KSHV infection, were grown in DMEM containing 10% FBS and 1% penicillin-streptomycin (Jones et al., 2012).

### Chemicals

*S*-(5′-adenosyl)-L-methionine chloride dihydrochloride (SAM) (A7007), N-acetyl-L-cysteine (NAC) (A9165), and cycloleucine (A48105) were purchased from Millipore Sigma.

### siRNA knockdown

iSLK-RGB-BAC16 cells seeded in 6-well plates at 50% confluency for 24 h were transfected with siRNAs with lipofectamine RNAiMAX (Thermo Fisher Scientific, 13778150). The knockdown efficiency was confirmed by reverse transcription real-time quantitative PCR (RT-qPCR) and Western-blotting. The sequences of the siRNAs targeting METTL16 were 5’-AACCAAAUUUUCAGGCUUG-3’ (MilliporeSigma, si1SASI_Hs02_00356556) and 5’-AAGGGAAGAUUUUGGACUU-3’ (MilliporeSigma, SASI_Hs01_00067253). The sequences of the siRNAs targeting MAT2A were 5’- CUGAUGCCAAAGUAGCUUG-3’ (Millipore Sigma, SASI_Hs01_00041621) and 5’- CCCAGAUAAGAUUUGUGAC-3’ (Millipore Sigma, SASI_Hs01_00041617), respectively. Universal negative control 1 (MilliporeSigma, SIC001) was used as a control siRNA.

### Plasmids

The METTL16 expression construct was generated by amplifying the coding sequence of METTL16 using cDNA prepared from iSLK cells as the template. The amplified fragment was cloned into the NotI/BamHI restriction enzyme sites of the PITA- Puro lentiviral vector with a flag-tag. The constructed plasmid was verified by DNA sequencing. The METTL16 plasmid DNA was transfected into cells using Lipofectamine 2000 according to the instructions of the manufacturer (ThermoFisher, 11668019).

### Virus titration

iSLK-RGB-BAC16 cells were induced into lytic replication under the specified conditions as previously described (Meng and Gao, 2021). At day 5 post-induction, the supernatant of iSLK-RGB-BAC16 cells was collected and used to infect MM cells by two-fold serial dilution in the presence of 10 μg/ml polybrene. MM cells were subjected to centrifugation at 1,500 rpm for 1 h after addition of the supernatant. The numbers of cells expressing mRFP1 were counted and photographed using an inverted fluorescence microscope at 48 h post-infection (hpi).

### Reverse transcription real-time quantitative PCR (RT-qPCR)

Total RNA was harvested with the TRIzol™ Reagent (ThermoFisher, T9424) according to the manufacturer’s instructions. The reverse transcription was carried out with 1 μg total RNA using the Maxima Hminus First Strand cDNA Synthesis Kit (Thermo Fisher, K1652). Quantitative PCR (qPCR) was carried out with the SsoAdvanced Universal SYBR Green Supermix Kit (Bio-Rad, 172-5272) using a CFX Connect Real-Time PCR Detection System (BioRad) according to the instructions of the manufacturer. The GAPDH was utilized as an internal control for normalization. The primers used are: CAATGACCCCTTCATTGACC (forward) and GATCTCGCTCCTGGAAGATG (reverse) for GAPDH; TCTGATCGGTCACCAGGATT (forward) and AAGTTATTCTGTTCCACATT (reverse) for METTL16; TTCTTGTTCAGGTCTCTTAT (forward) and TGACTAGCAACAGGCTTTAC (reverse) for MAT2A retained intron; TTCTTGTTCAGGTCTCTTAT (forward) and TATCCAGATCCCTGACAATGAC (reverse) for MAT2A mRNA; CAGACTTGTTGGCGTAGGCT (forward) and CCTTTCCCTCAGAGCTTGAA (reverse) for MAT2A m^6^A1; TTCTGAACAGCTGGTGTAGC (forward) and CTCTCTGAAGGAAGCTGGT (reverse) for MAT2A m^6^A2-6; GCTGCACATCAAGGTGCTAA (forward) and GCAACGTTCTGCAGTTCACA (reverse) for DICER m^6^A peak; CGATGCTGGCGCTGAGTACG (forward) and CATGGTGGTGAAGACGCCAG (reverse) for GAPDH as m^6^A peak normalization; CACAAAAATGGCGCAAGATGA (forward) and TGGTAGAGTTGGGCCTTCAGTT (reverse) for RTA; CGACAAAGTGAGGTGGCATTT (forward) and CCGCACACCACTTTAGTCCAA (reverse) for PAN RNA; AGGTCCCCCTCACCAGTAAA (forward) and GAGGACGTGTGTTTTGACCG (reverse) for ORF57; ATATGTCGCAGGCCGAATAC (forward) and CCACCCATCCTCCTCAGATA (reverse) for ORF65; and CATGCTGATGCGAATGTGC (forward) and AGCTTCAACATGGTGGGAGTG (reverse) for ORF-K8.

### Methylated RNA immunoprecipitation (MeRIP) RT-qPCR

MeRIP-RT-qPCR was performed as previously described with minor modifications (Tan et al., 2018). Briefly, the total RNA was fragmented with the RNA Fragmentation Reagents Kit (Thermo Fisher, AM8740) according to the manufacturer’s instructions. The sizes of RNA fragments were verified to be around 100 nucleotides using an Agilent 2100 Bioanalyzer System. Fragmented RNA at 6 μg were added to 10 μl Slurry of Pierce^TM^ Protein A Agarose beads pre-incubated with 1 μg of an anti-m^6^A polyclonal antibody (Synaptic Systems, 202-003) by rocking at 4°C for 3 h. The beads were washed 7 times with ice cold PBS. The beads were incubated with elution buffer at pH 7.4 containing 200 mM Tris-HCl, 1 M NaCl, 1% Igepal CA-630, and 20 mM N^6^- methyladenosine 5′-monophosphate sodium salt for 1 h at 4°C. Eluate was collected and subjected to GeneJET RNA Cleanup and Concentration Micro Kit for RNA purification (Thermo Fisher, K0841). Total RNA at 1 μg was used as the input. All eluted RNA were used for reverse transcription.

### RNA pull-down

METTL16 overexpressed iSLK-RGB-BAC16 cells were collected and lysed for 2 h at 4°C in immunoprecipitation (IP) buffer at pH 7.4 containing 250 mM Tri-HCl, 150 mM NaCl, 1 mM EDTA, 1% Igepal 630 and RNase inhibitor. The lysate was incubated with M2 anti-flag beads (Sigma, A2220) for 12 h at 4°C with gentle rocking. Beads were washed with cold IP buffer for 3 times. The RNA was extracted from beads using the TRIzol™ Reagent (ThermoFisher).

### Western-blotting

Cells were washed with ice cold PBS and then lysed in the sample buffer. The proteins were separated with an ExpressPlus^TM^ PAGE gel (Genscript, M42012) and transferred to a nitrocellulose membrane (GE Healthcare, 10600004). Following blocking with 5% milk for 1 h at room temperature, the membrane was incubated with a primary antibody at 4°C overnight and then probed with a secondary antibody conjugated with horseradish peroxidase (HRP). The signal was revealed using the SuperSignal West Femto Maximum Sensitivity Substrate (Thermo Fisher, 34096) and imaged with a ChemiDoc MP Imaging System (Bio-Rad, 17001402).

### Flow cytometry

The lytic cells were detected based on the expression of EGFP. Briefly, 1x10^5^ cells were collected by trypsinization and washed with PBS. The cells were examined with a BD LSRFortessa^TM^ Flow Cytometry System and the data were analyzed using the FlowJo software. To detect intracellular ROS, the CellROX Deep Red Reagent (Thermo Fisher, C10422) was added to the cells and incubated for 30 min. Cells were then collected for flow cytometry analysis.

### Analysis of metabolites by mass spectrometry

Cells (2×10^6^) cultured in 6 well plates were depleted of medium by aspiration and washed with 1 mL ice-cold 150 mM ammonium acetate (NH_4_CH_3_CO_2_) at pH 7.3. One mL of -80°C cold 80% MeOH was added. The samples were incubated at -80°C for 20 min. Cells were scraped off and pelleted at 4°C for 5 min at 1,500 *rpm*. The supernatant was transferred into new microfuge tubes and dried at room temperature under vacuum, and the metabolites were resuspended in water for LC-MS analysis.

Samples were randomized and analyzed on a Q-Exactive Plus hybrid quadrupole-Orbitrap mass spectrometer coupled to Vanquish UHPLC system (ThermoFisher). The mass spectrometer was run in polarity switching mode (+3.00 kV/- 2.25 kV) with an m/z window ranging from 65 to 975. Mobile phase A was 5 mM NH_4_AcO, pH 9.9, and mobile phase B was acetonitrile. Metabolites were separated on a Luna 3 μm NH2 100 NH2 100Å (150 x 2.0 mm) column (Phenomenex). The flow rate was 0.3 mL/min, and the gradient was from 15% A to 95% A in 18 min, followed by an isocratic step for 9 min and re-equilibration for 7 min. All samples were run in biological triplicate. Metabolites were detected and quantified as area under the curve based on retention time and accurate mass (5 ppm) using the TraceFinder 4.1 software (ThermoFisher).

### Statistical analysis

Results were presented as means plus standard deviations. All statistical analysis was done with PRISM 5.01 (GraphPad Software, USA) using 2-tailed Student’s t test or Two-way ANOVA followed by Tukey post hoc test if multiple samples were analyzed. Each experiment was independently repeated at least three times.

## Results

### METTL16 suppresses KSHV lytic replication

To determine the role of METTL16 in KSHV life cycle, we performed siRNA-mediated knockdown of METTL16 in iSLK-RGB-BAC16 cells (Fig. 1A). RGB-BAC16 is a recombinant KSHV expressing a monomeric red fluorescent protein 1 (mRFP1) under the control of a constitutively active elongation factor 1 promoter and an enhanced green fluorescent protein (EGFP) under the control of KSHV lytic PAN promoter for tracking KSHV latent and lytic replication phases, respectively (Brulois et al., 2014). Fluorescent microscopy examination revealed that all cells expressed mRFP1 but only few cells expressed EGFP under uninduced condition, indicating that KSHV was in tight latency in this system (Fig. 1B). METTL16 knockdown did not obviously alter the expression of mRFP1 or EGFP, indicating that METTL16 knockdown was insufficient to disrupt KSHV latency in most cells (Fig. 1B). Sodium butyrate (NaB), a histone deacetylase inhibitor, is a potent inducer of KSHV lytic replication (Miller et al., 1996). NaB treatment induced EGFP expression in close to 20% of the cells (Fig. 1B). METTL16 knockdown more than doubled the cells expressing EGFP (Fig. 1B). As expected, NaB treatment also increased the intensity of mRFP1 expression in cells, which was not affected by METTL16 knockdown. To further confirm these results, we analyzed the cells by flow cytometry. In agreement with the results observed by fluorescent microscope examination, NaB induced EGFP expression in 19% of the cells, which were further increased to 32-56% following METTL16 knockdown (Fig. 1C). Under uninduced condition, less than 0.3% of the cells expressed EGFP with or without METTL16 knockdown (Fig. 1C).

**Figure 1.**
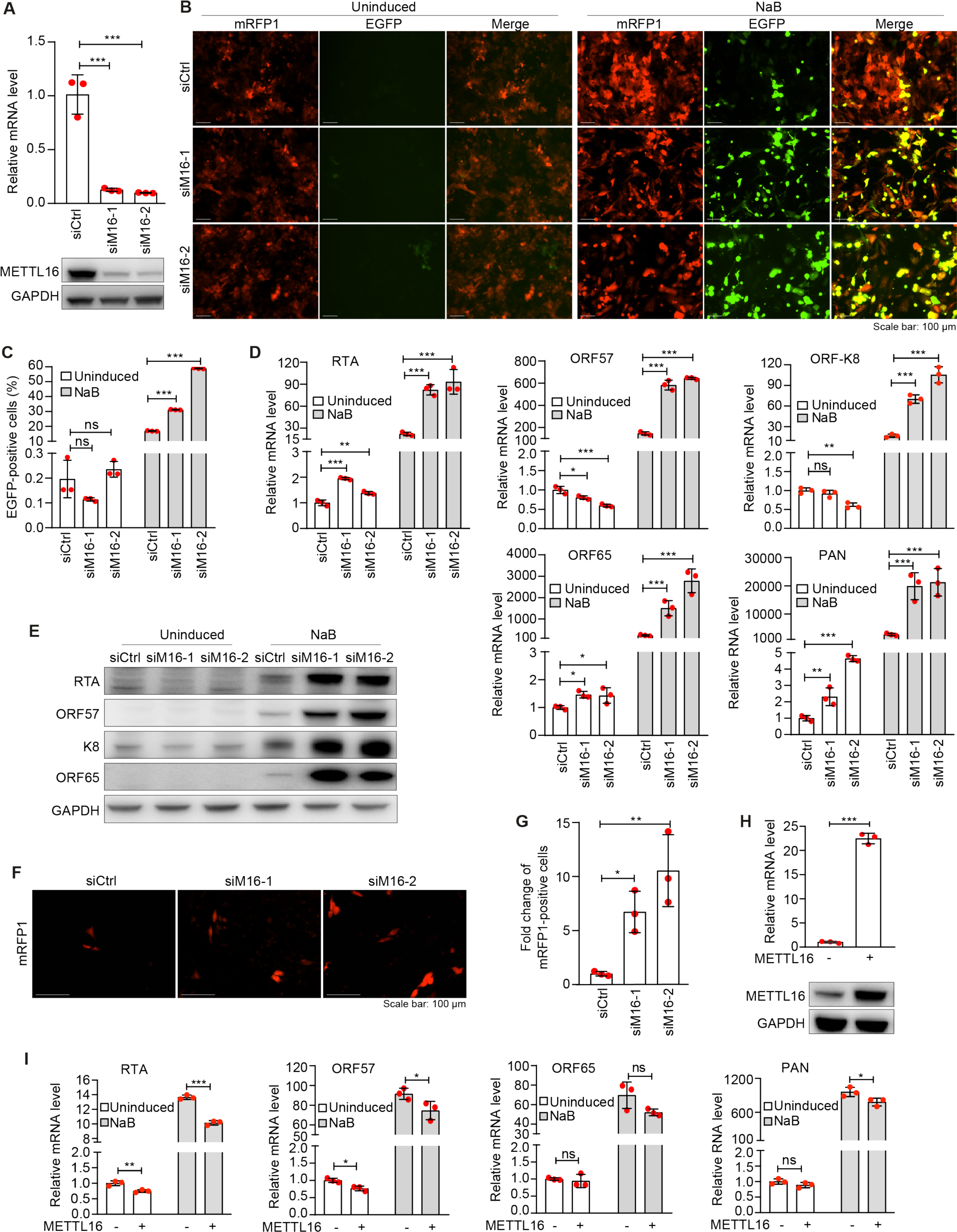
METTL16 suppresses KSHV lytic replication. **(A)** iSLK-RGB-BAC16 cells were transfected with siRNAs targeting METTL16 (siM16-1 and siM16-2) or the control siRNA (siCtrl) for 48 h. RNA or protein were harvested, and analyzed by RT-qPCR or Western-blotting to monitor the knockdown efficiency. GAPDH was used as normalization control. **(B, C)** iSLK-RGB-BAC16 cells were transfected with siRNAs targeting METTL16 or the control siRNA for 24 h. Cells were treated with 3 mM of sodium butyrate (NaB) for 48 h and observed with a fluorescent microscope **(B)** and analyzed by flow cytometry for EGFP-positive cells **(C)**. **(D)** iSLK-RGB-BAC16 cells transfected with siRNAs targeting METTL16 or the control siRNA were treated with 3 mM of NaB for 72 h and examined for the expression level of KSHV transcripts RTA, ORF57, ORF-K8 and ORF65 by RT-qPCR. GAPDH was used as normalization control. **(E)** iSLK-RGB-BAC16 cells transfected with siRNAs targeting METTL16 or the control siRNA were treated with 3 mM of NaB for 72 h. Cells were lysed and analyzed for the expression of RTA, ORF57, K8, and ORF65 proteins by Western-blotting. GAPDH was used as a loading control. **(F, G)** iSLK-RGB-BAC16 cells transfected with siRNAs targeting METTL16 or the control siRNA were treated with 3 mM of NaB for 48 h. The cells were then washed with PBS and cultured in fresh medium without NaB, hygromycin, puromycin and G418 for 72 h. Supernatants were collected and used to infect MM cells. mRFP1-positive cells were imaged **(F)** and quantified at 48 h post- infection **(G)**. **(H)** iSLK-RGB-BAC16 cells were transfected with a plasmid expressing METTL16 or a vector control. RNA and protein were harvested and analyzed by RT- qPCR and Western-blotting to monitor the overexpression efficiency. GAPDH was used as a normalization control. **(I)** iSLK-RGB-BAC16 cells with overexpression of METTL16 were treated with or without 3 mM of NaB for 72 h. Levels of KSHV transcripts RTA, ORF57, PAN RNA and ORF65 were analyzed by RT-qPCR. *, **, and *** indicate *p* values of <0.05, <0.01, and <0.001, respectively, ns, not significant.

We next determined whether METTL16 knockdown could affect KSHV lytic transcriptional program. Under uninduced condition, METTL16 knockdown increased the expression of KSHV lytic transcripts RTA, ORF65 and PAN RNA while slightly decreased the expression of ORF57 and ORF-K8 transcripts (Fig. 1D). However, these effects were marginal, which were consistent with the results of EGFP expression (Fig 1B and 1C) and the expression of KSHV lytic proteins RTA, ORF57, ORF-K8 and ORF65 revealed by Western-blotting (Fig 1E). NaB treatment significantly increased the expression of KSHV lytic transcripts ranging from 22- to 2,000-fold while METTL16 knockdown further increased the expression of lytic transcripts by 4.1- to 14-fold (Fig 1D), which were confirmed by Western-blotting (Fig 1E).

To determine whether METTL16 knockdown promoted the production of infectious KSHV virions, culture supernatants were collected at day 5 post-induction with NaB and used to infect MM cells, which are highly susceptible to KSHV infection. The numbers of mRFP1-positive cells at day 2 post-infection were examined to determine the relative titers of KSHV infectious virions produced by the cultures. METTL16 knockdown increased the production of infectious virions by 7- to 10-fold (Fig. 1F and 1G).

To confirm the role of METTL16 in KSHV life cycle, we overexpressed METT16 in iSLK-RGB-BAC16 cells (Fig. 1H), and examined the expression of KSHV lytic genes by RT-qPCR. METTL16 overexpression significantly inhibited the expression of KSHV lytic transcripts under NaB induction (Fig. 1I) but the degrees of change were less than those observed after METTL16 knockdown, which could be due to the presence of endogenous expression of METTL16. In uninduced cells, KSHV lytic transcripts were also slightly reduced (Fig. 1I).

Taken together, these results indicated that METTL16 had a suppressive role in KSHV lytic replication program. METTL16 knockdown enhanced while overexpression inhibited KSHV lytic replication.

### METTL16 regulates the expression of MAT2A

Previous studies have shown that METTL16 binding to and methylation of the hairpin sites in the MAT2A 3’UTR regulate the maturation and degradation of its transcript (Pendleton et al., 2017). At a low SAM level, METTL16 occupancy at hp1 site promotes MAT2A transcript splicing and maturation while at a high SAM level, METTL16 methylation of hairpin sites promotes MAT2A transcript degradation (Fig. 2A). To determine whether METTL16 methylated MAT2A 3’UTR in iSLK-RGB-BAC16 cells, we performed MeRIP-RT-qPCR following METTL16 knockdown. METTL3-mediated m^6^A methylation of Dicer transcript was used as a negative control. METTL16 knockdown significantly decreased the methylation of MAT2A transcript at 1^st^ site and 2^nd^-6^th^ sites but had no effect on the DICER m^6^A site (Fig. 2B). As a result, METTL16 knockdown significantly increased the level of MAT2A transcript with retained intron but reduced the level of mature MAT2A transcript and MAT2A protein (Fig. 2C and 2D). In contrast, METTL16 overexpression upregulated the methylation levels at MAT2A 1^st^ and 2^nd^-6^th^ sites (Fig. 2E). As a result, METTL16 overexpression increased the level of mature MAT2A transcript (Fig. 2F). Furthermore, RNA pull-down assay indeed showed METTL16 binding to the MAT2A transcript (Fig. 2G).

**Figure 2.**
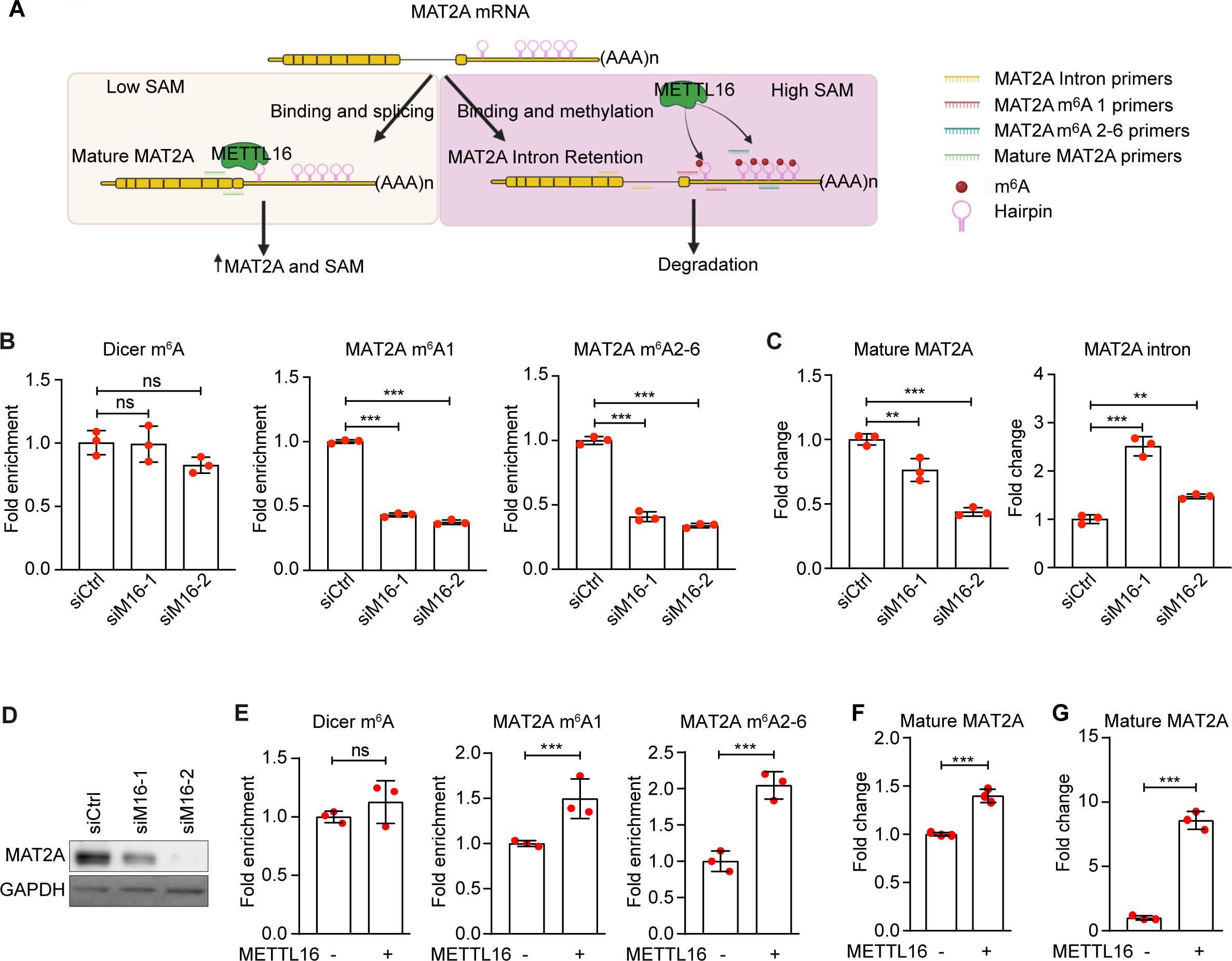
METTL16 binds and methylates MAT2A to mediate MAT2A splicing and expression. **(A)** Schematic illustration of METTL16 regulation of MAT2A expression in iSLK-RGB-BAC16 cells. **(B**) iSLK-RGB-BAC16 cells were transfected with siRNAs targeting METTL16 (siM16) or the control siRNA (siCtrl), and collected for RNA purification. The level of m^6^A on a specific transcript locus was examined by MeRIP- qPCR. m^6^A peaks located at MAT2A hp1 site and hp2-6 sites were detected. Dicer m^6^A peak was used as a negative control. **(C)** iSLK-RGB-BAC16 cells were transfected with siRNAs targeting METTL16 or the control siRNA and collected for RNA purification. The expression levels of mature MAT2A and retained intron transcripts were examined by qRT-PCR. GAPDH was used as a normalization control. **(D)** iSLK-RGB-BAC16 cells were transfected with siRNAs targeting METTL16 or the control siRNA. Cells were lysed and the expression of MAT2A protein was detected by Western-blotting. GAPDH was used as a loading control. **(E)** iSLK-RGB-BAC16 cells transfected with a plasmid expressing METTL16 or a vector control were collected for RNA purification. The level of m^6^A on a specific transcript locus was examined by MeRIP-qPCR. m^6^A peaks located at the MAT2A hp1 site and hp2-6 sites were detected. Dicer m^6^A peak was used as a negative control. **(F)** The expression level of mature MAT2A in METTL16 overexpressed iSLK-RGB-BAC16 cells were detected by RT-qPCR. **(G)** iSLK-RGB-BAC16 cells transfected with a plasmid expressing METTL16 or a vector control were collected for RNA purification. RNA immunoprecipitation (RIP)-qPCR was performed and fold of enrichment of MAT2A RNA was calculated. *, **, and *** indicate *p* values of <0.05, <0.01, and <0.001, respectively, ns, not significant.

These results collectively indicated that METTL16 bound to MAT2A transcript at 3’UTR, and promoted its splicing, maturation and expression in iSLK-RGB-BAC16 cells (Fig. 2A).

### MAT2A suppresses KSHV lytic replication and mediates METTL16 regulation of KSHV lytic replication

To explore whether MAT2A could regulate KSHV lytic replication, we performed siRNA-mediated MAT2A knockdown in iSLK-RGB-BAC16 cells with or without NaB induction (Fig. 3A). Examination by fluorescence microscopy revealed that, similar to METTl16 knockdown, MAT2A knockdown increased the number of EGFP-positive cells in the presence of NaB induction but only had marginal effect on the uninduced cells (Fig. 3B). Flow cytometry analysis showed that MAT2A knockdown increased the number of EGFP-positive cells from 12% to 22-42% in NaB-induced cells (Fig. 3C) and slightly increased of EGFP-positive cells from 1.2% to 2.6-3.8% in uninduced cells (Fig. 3C). Examination of KSHV lytic proteins including RTA, ORF57, ORF-K8 and ORF65 by Western-blotting further confirmed that MAT2A knockdown increased the expression of these viral lytic proteins in NaB-induced cells while had minimal effect on the uninduced cells (Fig. 3D). These results indicated that MAT2A suppressed KSHV lytic replication.

**Figure 3.**
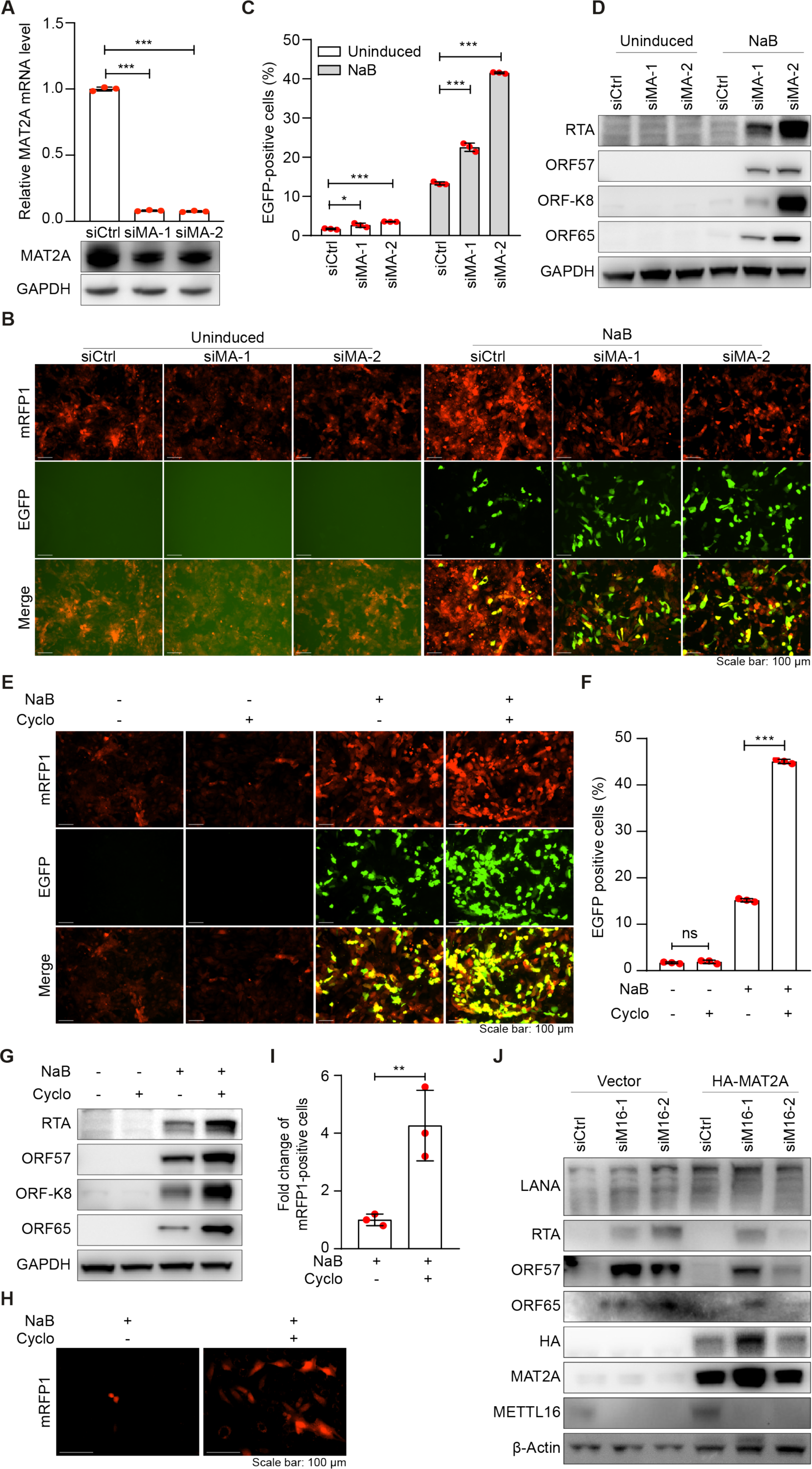
Inhibition of MAT2A promotes KSHV lytic replication. **(A)** iSLK-RGB- BAC16 cells were transfected with siRNAs targeting MAT2A (siMA-1 and siMA-2) or the control siRNA (siCtrl). MAT2A mRNA and protein levels were analyzed by RT-qPCR and Western-blotting. GAPDH was used as a normalization control. **(B, C)** iSLK-RGB- BAC16 cells were transfected with siRNAs targeting MAT2A or the control siRNA for 24 h. Cells were treated with 3 mM of sodium butyrate (NaB) for 48 h and observed with a fluorescent microscope **(B)** and analyzed by flow cytometry for EGFP-positive cells **(C)**. **(D)** iSLK-RGB-BAC16 cells transfected with siRNAs targeting MAT2A or the control siRNA were treated with 3 mM of NaB for 72 h and examined for the expression of KSHV proteins RTA, ORF57, ORF-K8 and ORF65 by Western-blotting. GAPDH was used as a loading control. **(E, F)** iSLK-RGB-BAC16 cells were treated with 50 mM of cycloleucine (Cyclo) and 3 mM of NaB for 48 h and observed with a fluorescent microscope **(E)** and analyzed by flow cytometry for EGFP-positive cells **(F)**. **(G)** iSLK- RGB-BAC16 cells treated with 50 mM of cycloleucine (Cyclo) and 3 mM of NaB for 72 h and examined for the expression of KSHV lytic proteins by Western-blotting. GAPDH was used as a loading control. **(H, I)** iSLK-RGB-BAC16 cells were treated with 50 mM of cycloleucine (Cyclo) and 3 mM of NaB for 24 h. Cells were then washed with PBS and cultured in fresh medium without NaB, hygromycin, puromycin and G418 for 72 h. Supernatants were collected and used for titrating infectious virions by infecting MM cells. mRFP1-positive cells were imaged **(H)** and quantified at 48 h post-infection **(I)**. **(J)** iSLK-RGB-BAC16 cells were transfected with siRNAs targeting MAT2A or the control siRNA. After 24 h, cells were transfected with a plasmids expressing HA-tagged MAT2A (HA-MAT2A) or a vector control (Vector) and treated with 3 mM of NaB. Cells were lysed to collect the proteins at 72 h post-treatment. Antibodies to HA, MAT2A, METTL16 and KSHV proteins LANA, RTA, ORF57 and ORF65 were used to detect their respective proteins by Western-blotting. β-actin was used as a loading control. *, **, and *** indicate *p* values of <0.05, <0.01, and <0.001, respectively, ns, not significant.

To confirm the role of MAT2A on KSHV lytic replication, we treated the cells with cycloleucine, a substrate-competitive inhibitor with a low affinity to MAT2A (Quinlan et al., 2017). Similar to siRNA-mediated MAT2A knockdown, cycloleucine treatment increased the numbers of EGFP-positive cells upon NaB induction but had minimal effect on uninduced cells (Fig. 3E and 3F). In agreement with these results, cycloleucine treatment increased the levels of KSHV lytic proteins including RTA, ORF57, ORF-K8 and ORF65 in NaB-induced cells but had minimal effect on the uninduced cells (Fig. 3G). Furthermore, Cycloleucine treatment significantly increased the yield of infectious virions by 4 folds in NaB-induced cells (Fig. 3H and 3I). Taken together, these results demonstrated that pharmacological inhibition of MAT2A promoted KSHV lytic replication but had minimal effect on KSHV latency.

In addition to MAT2A, METTL16 might also regulate the expression and functions of other cellular genes. To determine whether METTL16 regulated KSHV replication through MAT2A, we performed a rescue experiment by expressing MAT2A in METTL16 knockdown iSLK-RGB-BAC16 cells. Overexpression of MAT2A partially inhibited the increased expression of KSHV lytic proteins RTA, ORF57 and ORF65 caused by METTL16 knockdown (Fig. 3J). Hence, METTL16 controlled KSHV lytic replication at least in part by binding to MAT2A transcript, and regulating its maturation and expression.

### METTL16 inhibits the KSHV lytic replication by regulating intracellular SAM

MAT2A is a key regulator of methionine-SAM cycle by catalyzing the conversion of L-methionine to SAM in the presence of adenosine triphosphate (ATP) (Bottiglieri, 2002). SAM can be converted to *S*-adenosyl-L-homocysteine (SAH) in the methionine cycle or polyamines, or used as a methyl donor in the methylation of protein, DNA and RNA (Fig. 4A). We determined the intracellular SAM level by mass spectrometry following METTL16 or MAT2A knockdown. As expected MAT2A knockdown significantly decreased the intracellular SAM level in both uninduced and NaB-induced cells, leading to the lower level of SAH, a product of downstream of SAM (Fig. 4B).

**Figure 4.**
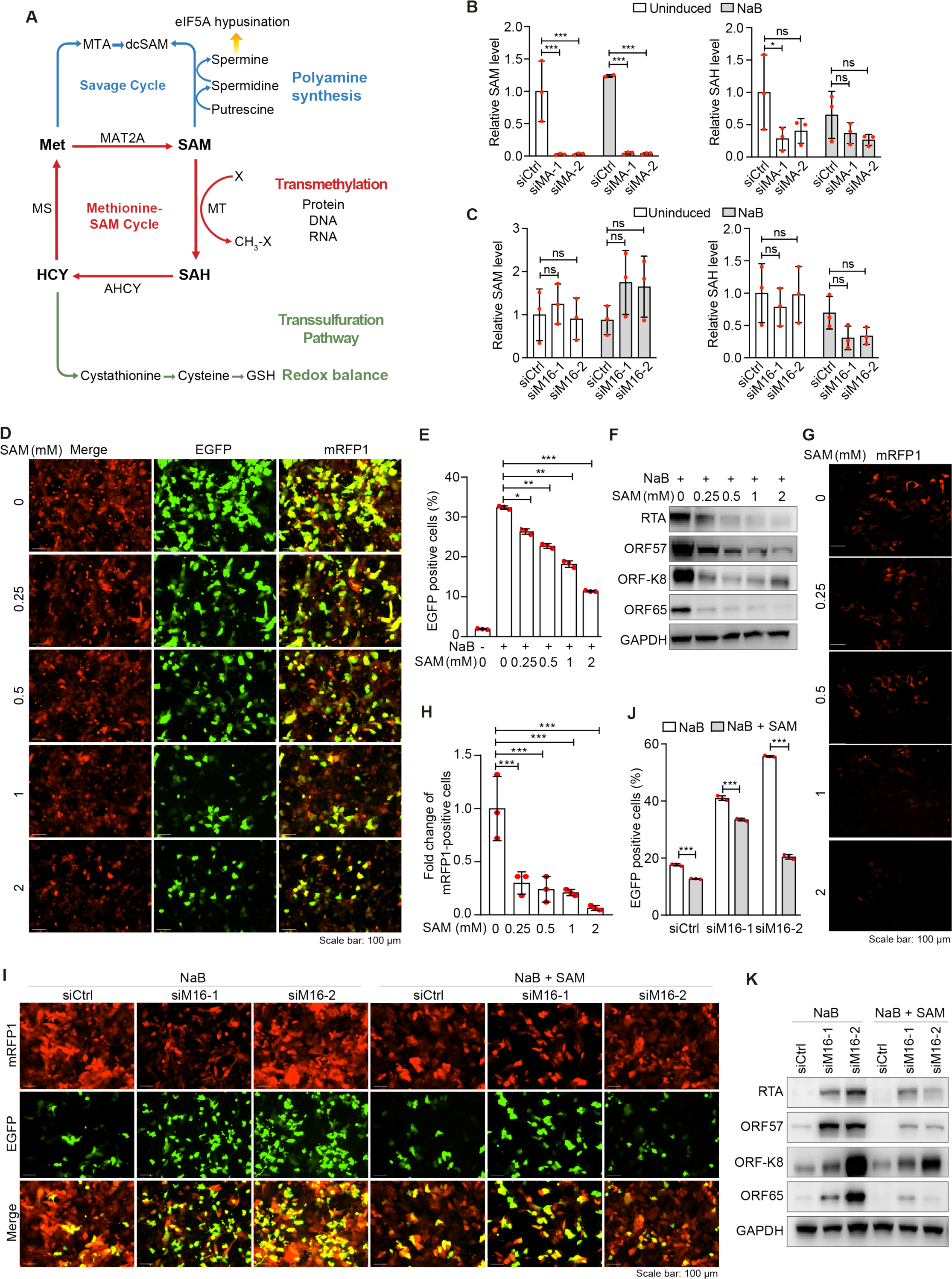
METTL16 and MAT2A mediates the KSHV lytic replication by regulating intracellular SAM level. **(A)** Schematic illustration of methionine-SAM cycle and downstream pathways. **(B)** iSLK-RGB-BAC16 cells were transfected with siRNAs targeting MAT2A (siMA-1 and siMA-2) or the control siRNA (siCtrl). Cells were then treated with 3 mM of NaB for 24 h, and analyzed for SAM and SAH levels by mass spectrometry. **(C)** iSLK-RGB-BAC16 cells were transfected with siRNAs targeting METTL16 (siM16-1 and siM16-2) or the control siRNA (siCtrl). Cells were then treated with 3 mM of NaB for 24 h, and analyzed for SAM and SAH levels by mass spectrometry. **(D, E)** iSLK-RGB-BAC16 cells induced with 3 mM of NaB were treated with 0, 0.25, 0.5, 1 or 2 mM of SAM for 48 h, and observed with a fluorescent microscope **(D)** and analyzed by flow cytometry for EGFP-positive cells **(E)**. **(F)** iSLK- RGB-BAC16 cells induced with 3 mM NaB were treated with SAM at the indicated concentrations for 72 h. Cells were lysed and KSHV lytic proteins were analyzed by Western-blotting. GAPDH was used as a loading control. (**G, H)** iSLK-RGB-BAC16 cells induced with 3 mM NaB were treated with SAM at the indicated concentrations for 24 h. Cells were then washed with PBS and cultured in fresh medium without NaB, hygromycin, puromycin and G418 for 72 h. Supernatants were collected and used for titrating infectious virions by infecting MM cells. mRFP1-positive cells were imaged **(G)** and quantified at 48 h post-infection **(H)**. **(I, J)** iSLK-RGB-BAC16 cells were transfected with siRNAs targeting METTL16 or the control siRNA for 24 h. Cells were then treated with 3 mM of NaB and 1 mM of SAM for 72 h, and observed with a fluorescent microscope **(I)** and analyzed by flow cytometry for EGFP-positive cells **(J)**. **(K)** iSLK-RGB-BAC16 cells transfected with siRNAs targeting METTL16 or the control siRNA were treated with 3 mM of NaB and 1 mM of SAM for 72 h and collected. KSHV lytic proteins were analyzed by Western-blotting. GAPDH was used as a loading control. *, **, and *** indicate *p* values of <0.05, <0.01, and <0.001, respectively, ns, not significant.

METTL16 knockdown also decreased the level of SAH in NaB-induced cells but not in uninduced cells. However, no significant change was observed for SAM level in uninduced and NaB-induced cells (Fig. 4C). These results suggested that METTL16 might have additional targets besides MAT2A or that the reduced SAM level as a result of lower MAT2A expression might be compensated by slower conversion of SAM to other downstream products or pathways including polyamines, and methylation of protein, DNA or RNA (Fig. 4A). The fact that SAH level was decreased following METTL16 knockdown in NaB-induced cells suggested that the downstream products of SAH including those in the methionine cycle such as homocysteine (HCY) and the transsulfation pathway such as cysthionine, cysteine and glutathione might also be affected (Fig. 4A). Since numerous downstream pathways of SAM including transamination (polyamine synthesis), transmethylation (methylation of protein, DNA and RNA) and transsulfuration (glutathione synthesis) have been implicated in KSHV lytic replication (Choi et al., 2022; Fiches et al., 2022; Gao et al., 2019; Gunther and Grundhoff, 2010; Li and Gao, 2023; Tan et al., 2018; Toth et al., 2010; Ye et al., 2011b), we further examined the role of SAM in the KSHV lytic replication. SAM treatment reduced the numbers of GFP-positive cells in a concentration-dependent manner in NaB-induced cells (Fig. 4D and 4E). In agreement with these results, SAM treatment decreased the levels of KSHV lytic proteins including RTA, ORF57, ORF-K8 and ORF65 (Fig. 4F). Furthermore, SAM treatment dramatically decreased the production of infectious virions, reaching 95% of inhibition at the highest dose of 2 mM (Fig. 4G and 4H).

To examine whether SAM could reverse the effect of METTL16 knockdown, iSLK-RGB-BAC16 cells were treated with 1 mM SAM for 2 h prior to NaB induction. SAM treatment reversed the effect of METTL16 knockdown on KSHV lytic replication (Fig. 4I and 4J). SAM treatment also inhibited KSHV lytic replication in cells transduced with the scramble control (Fig. 4I and 4J). The results were further confirmed by Western-blotting, which examined the expression levels of KSHV lytic proteins including RTA, ORF57, ORF-K8 and ORF65 (Fig. 4K).

Together, these results indicated that SAM suppressed KSHV lytic replication and reversed the enhanced effect of METTL16 knockdown on KSHV lytic replication. Thus, METLL16 might regulate SAM cycle to inhibit KSHV lytic replication by modulating the intracellular SAM availability.

### METTL16 controls oxidative stress to regulate KSHV lytic replication

SAM serves as methyl donor in almost all cellular methyl transfer reactions, which is critical for a variety of biological processes (Mentch and Locasale, 2016). Our results showed that MAT2A knockdown decreased both SAM and SAH while METTL16 knockdown decreased SAH but not SAM level in NaB-induced cells (Fig. 4B and 4C). SAH is the precursor of the major endogenous antioxidant glutathione (GSH) and regulates cellular redox homeostasis and ROS level (Serefidou et al., 2019; Tatekawa et al., 2021). Previous studies have shown that ROS is essential and sufficient for activating KSHV lytic replication and inducing the expression of KSHV lytic genes (Gao et al., 2019; Li and Gao, 2023; Ye et al., 2011b). Therefore, we examined the intracellular GSH level. MAT2A knockdown reduced the intracellular GSH level in both uninduced and NaB-induced cells (Fig. 5A). METTL16 knockdown appeared also to reduce GSH level with one of the siRNAs while the second one was inconsistent in uninduced cells (Fig. 5B). In agreement with the GSH results, METTL16 knockdown weakly increased the intracellular ROS level in uninduced cells (Fig. 5C). As previously reported (Gao et al., 2019), NaB treatment dramatically increased the intracellular ROS level, which was further increased following METTL16 knockdown (Fig. 5C). Consistently, MAT2A knockdown increased the intracellular ROS level in both uninduced and NaB-induced cells though there was no change with one of the siRNAs in NaB-induced cells (Fig. 5D). Furthermore, cycloleucine treatment increased while SAM treatment reduced the intracellular ROS level in both uninduced and NaB-induced cells (Fig. 5E and 5F).

**Figure 5.**
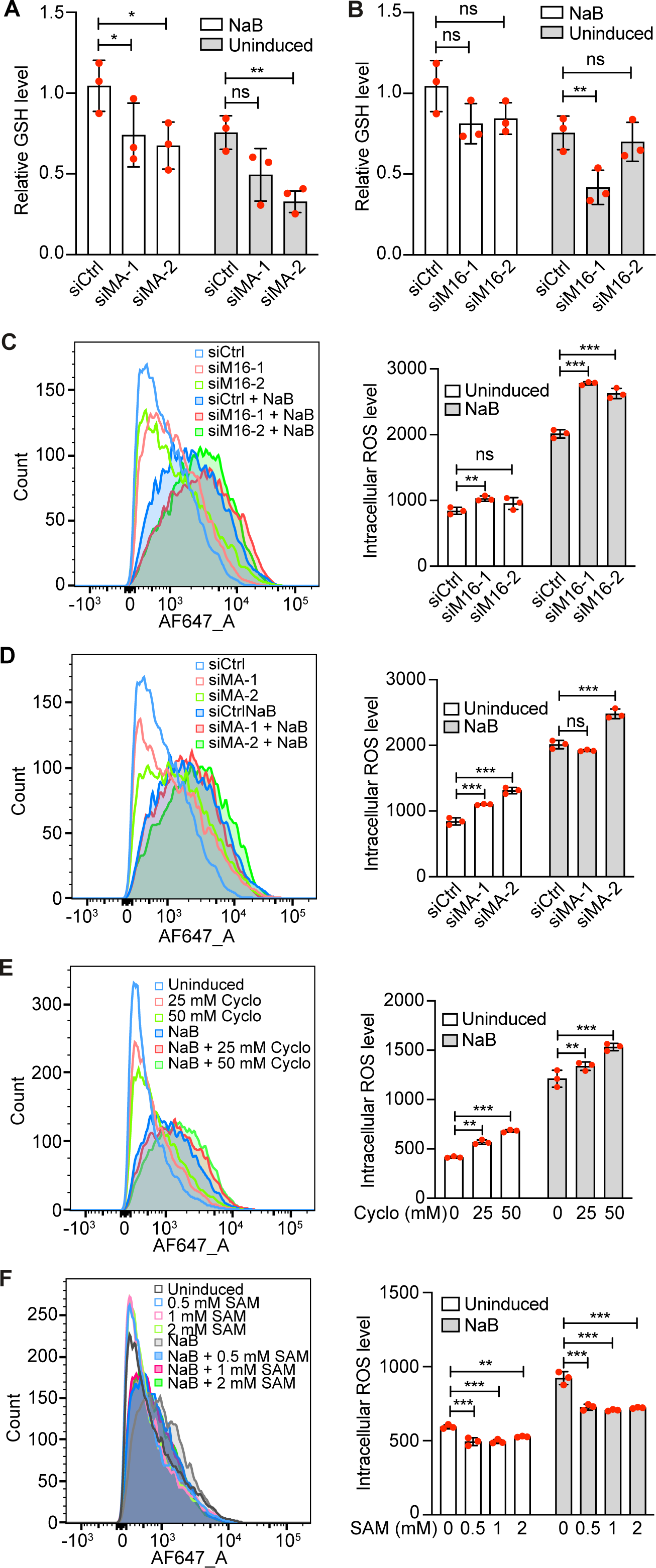
METTL16 and MAT2A regulate intracellular GSH and ROS levels. **(A)** iSLK-RGB-BAC16 cells transfected with siRNAs targeting MAT2A (siMA-1 and siMA-2) or the control siRNA (siCtrl) were treated with 3 mM of NaB for 24 h. The intracellular GSH level was analyzed by mass spectrometry. **(B)** iSLK-RGB-BAC16 cells transfected with siRNAs targeting METTL16 (siM16-1 and siM16-2) or the control siRNA (siCtrl) were treated with 3 mM of NaB for 24 h. The intracellular GSH level was analyzed by mass spectrometry. **(C)** iSLK-RGB-BAC16 cells transfected with siRNAs targeting MAT2A or the control siRNA were treated with 3 mM of NaB. The intracellular ROS level was measured and quantified by flow cytometry. **(D)** iSLK-RGB-BAC16 cells transfected with siRNAs targeting METTL16 or the control siRNA were treated with 3 mM of NaB. The intracellular ROS level was measured and quantified by flow cytometry. **(E)** ROS levels in iSLK-RGB-BAC16 cells treated with 3 mM of NaB and cycloleucine (Cyclo) at the indicated concentrations were measured by flow cytometry. **(F)** Intracellular ROS levels of iSLK-RGB-BAC16 cells treated with 3 mM of NaB and SAM at the indicated concentrations were measured by flow cytometry. *, **, and *** indicate *p* values of <0.05, <0.01, and <0.001, respectively, ns, not significant.

N-acetyl-L-cysteine (NAC) is a H_2_O_2_ scavenger that can effectively inhibit KSHV lytic replication (Gao et al., 2019; Ye et al., 2011b). Indeed, treatment with NAC slightly reduced the intracellular ROS level in METTL16 knockdown NaB-induced cells (Fig. 6A). Similarly, NAC also reduced the intracellular ROS level in MAT2A knockdown NaB- induced cells (Fig. 6B). In agreement with these results, NAC treatment reduced the numbers of EGFP-positive cells in METTL16 or MAT2A knockdown NaB-induced cells (Fig. 6C-F). These results further confirmed that METTL16 and MAT2A inhibited KSHV lytic replication by suppressing ROS and maintaining cellular redox homeostasis.

**Figure 6.**
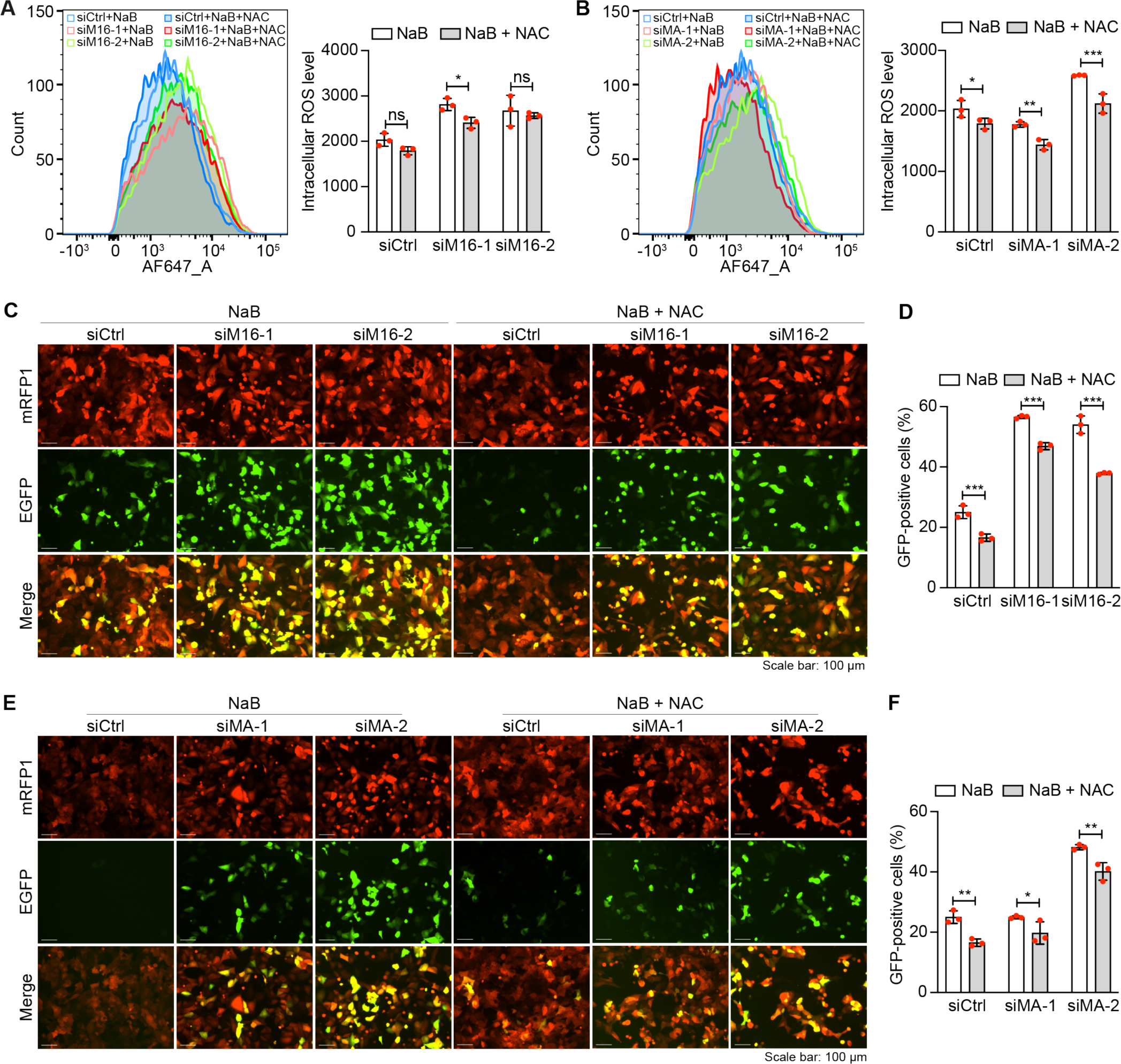
METTL16 and MAT2A regulate intracellular ROS level to control KSHV lytic replication. **(A)** iSLK-RGB-BAC16 cells pre-treated with NAC at 10 mM for 2 h were transfected with siRNAs targeting METTL16 (siM16-1 and siM16-2) or the control siRNA (siCtrl) for 24 h in the presence of NAC. Cells were then induced with 3 mM of NaB in the presence of NAC for 48 h and the intracellular ROS level was measured and quantified by flow cytometry. **(B)** iSLK-RGB-BAC16 cells pre-treated with NAC at 10 mM for 2 h were transfected with siRNAs targeting MAT2A (siMA-1 and siMA-2) or the control siRNA (siCtrl) for 24 h in the presence of NAC. Cells were then induced with 3 mM of NaB in the presence of NAC for 48 h and the intracellular ROS level was measured and quantified by flow cytometry. **(C, D)** iSLK-RGB-BAC16 cells pre-treated with NAC at 10 mM for 2 h were transfected with siRNAs targeting METTL16 or the control siRNA for 24 h in the presence of NAC and then induced with 3 mM of NaB. Cells were observed with a fluorescent microscope **(C)** and analyzed by flow cytometry for EGFP-positive cells **(D)**. **(E, F)** iSLK-RGB-BAC16 cells pre-treated with NAC at 10 mM for 2 h were transfected with siRNAs targeting MAT2A or the control siRNA for 24 h in the presence of NAC and then induced with 3 mM of NaB. Cells were observed with a fluorescent microscope **(E)** and analyzed by flow cytometry for EGFP-positive cells **(F)**.

## Discussion

As the most abundant internal mRNA modification, m^6^A is involved in all aspects of mRNA life cycle and regulates diverse cellular processes (Jiang et al., 2021; Zhang et al., 2021). Numerous viruses have been shown to utilize cellular m^6^A machinery to optimize their infections by either directly regulating the viral genomes or transcripts, or indirectly reprogramming the cellular epitranscriptome (Tan and Gao, 2018a, b). Recent studies have systematically profiled m^6^A modifications on KSHV RNA transcripts, which identified unique and conserved m^6^A modifications across different infection systems and viral replication phases, and elucidated the functions of m^6^A-related proteins including FTO, YTHDC1, YTHDF2, YTHDF3, METTL3, RBM15 and SND1 in KSHV life cycle (Baquero-Perez et al., 2019; Hesser et al., 2018; Martin et al., 2021; Tan et al., 2018; Ye et al., 2017). METTL3, the most well studied RNA methyltransferase, has been found to regulate KSHV life cycle in various manners depending on cell types and viral replication phases (Baquero-Perez et al., 2019; Hesser et al., 2018). YTHDF2 binds to KSHV transcripts and mediates their degradation (Tan et al., 2018) while YTHDC1 mediates the transcript splicing of ORF50, a transactivator that is necessary and sufficient for KSHV lytic replication (Ye et al., 2017). Besides, SND1, a newly proposed m^6^A reader protein, regulates ORF50 RNA stability to affect KSHV lytic replication (Baquero-Perez et al., 2019).

The role of METTL16, a newly identified RNA methyltransferase, in KSHV infection remains unclear. In this study, we have shown that siRNA-mediated METTL16 knockdown promotes KSHV lytic replication and enhances the expression of viral lytic genes. METTL16 knockdown also disrupted viral latency in a small number of KSHV- infected cells. These results indicate that METTL16 has a suppressive role in KSHV lytic replication.

METTL16 appears to bind to motifs different from those of METTL3 and methylate a smaller number of transcripts (Pendleton et al., 2017). Previous studies have found that METTL16 methylates U6 at A43 and 6 sites in the MAT2A 3’UTR (Pendleton et al., 2017). Under SAM-limiting conditions, the occupancy of METTL16 on MAT2A hp1 site is enhanced, leading to increased m^6^A writing and splicing of a MAT2A retained intron, and hence transcript maturation and expression. The enhanced expression of MAT2A increases the conversion of L-methionine to SAM. Thus, METTL16 acts in a positive feedback mechanism to ensure the replenishment of the low intracellular SAM level (Pendleton et al., 2017). At a high intracellular SAM level, METTL16 binding to and m^6^A writing of MAT2A transcript are reduced leading to inefficient transcript splicing and maturation as well as faster degradation (Pendleton et al., 2017). In this case, METTL16 functions in a negative feedback mechanism to control the high intracellular SAM level. Therefore, METTL16 is closely tied to the SAM cycle and behaves a “sensor” of SAM level not only functions as a “writer” but also as a “reader” to regulate MAT2A transcript splicing, maturation and stability. These dual METTL16 functions maintain intracellular SAM level and the homeostasis of the SAM cycle. In this study, we examined the regulation of MAT2A transcript by METTL16 in iSLK-RGB-BAC16 cells. We have found that METTL16 methylates MAT2A transcript hp1 and that METTL16 knockdown inhibits mature MAT2A transcript expression by decreasing splicing of its retained intron. More importantly, we have found that METLL16 regulation of KSHV lytic replication is tightly linked to the SAM cycle and the intracellular SAM level. Specifically, a higher intracellular SAM level inhibits KSHV lytic replication and controls viral latency while a lower intracellular SAM level promotes KSHV lytic replication.

In previous studies, METTL16-dependent peaks are not enriched at UACAGAGAA consensus which might be due to the presence of other cellular co- factors. However, based on the sites methylated by METTL16 on U6 and MAT2A, which were experimentally validated, the UACAGAGAA sequence might only partially reflect the requirements for METTL16 to carry out its function of methyltransferase (Doxtader et al., 2018; Pendleton et al., 2017). Since the lack of this consensus sequence on important KSHV transcripts, it is unlikely that METTL16 regulates KSHV lifecycle through direct methylation on the KSHV transcripts. In contrast, it is highly possible that METTL16 affects KSHV life cycle through regulating other cellular factors and pathways. In this study, we have shown that METTL16 mediates KSHV life cycle by regulating the splicing of MAT2A transcript and the intracellular SAM level. SAM is linked to multiple downstream pathways including polyamine synthesis, methylation of protein, DNA and RNA, and glutathione synthesis and ROS level (Giulidori et al., 1984; Kalhan and Marczewski, 2012), which have been linked to KSHV lytic replication (Choi et al., 2022; Fiches et al., 2022; Gao et al., 2019; Gunther and Grundhoff, 2010; Li and Gao, 2023; Tan et al., 2018; Toth et al., 2010; Ye et al., 2011b). We have found that METTL16 or MAT2A knockdown reduces intracellular SAM and SAH levels leading to a lower GSH level and a higher ROS level, and hence increased KSHV lytic replication. Thus, METTL16 regulates KSHV life cycle by connecting m^6^A epitranscriptome to cellular metabolism (Fig. 7). As METTL16 is a SAM “sensor”, it functions as a thermostat to promote KSHV latency by controlling the intracellular ROS level. Nevertheless, since the UACAGAGAA sequence only partially reflects the consensus motif of METTL16, it remains possible that METTL16 might promote KSHV lytic replication by directly methylating KSHV transcripts.

**Figure 7.**
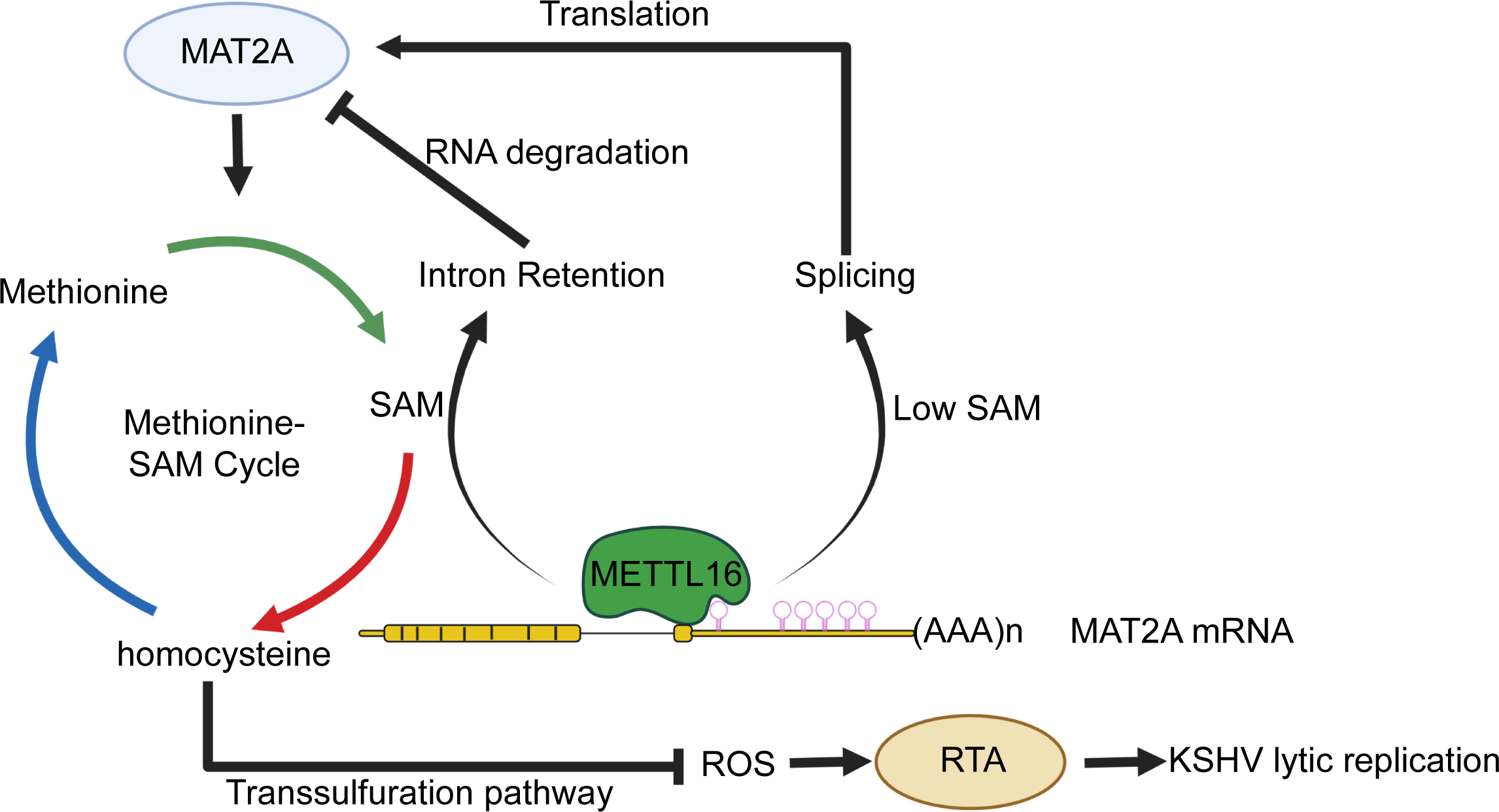
Schematic illustration of METTL16 mediating the KSHV lytic replication by regulation of cellular SAM level.

## Acknowledgments

We thank members of Drs. Shou-Jiang Gao’s lab for technical assistances and discussions. This study was supported by grants from the National Institutes of Health (CA096512 and CA124332 to S.-J. Gao) and in part by award P30CA047904.

## Author contributions

S.-J. Gao conceived, designed, supervised and managed the project. X. Q. Zhang, W. Meng, J. Feng, X. H. Gao and C. Qin performed the experiments. X. Q. Zhang, W. Meng, C. Qin, P. H. Feng, Y. F. Huang and S.-J. Gao interpreted and analyzed the data. X. Q. Zhang, W. Meng and S.-J. Gao wrote the manuscript with inputs from all the authors. All the authors read, reviewed and approved the manuscript.

## Data availability

The data that support the findings of this study are available from the corresponding author upon reasonable request.

